# Biomechanical effects of load-carrying in adolescent idiopathic scoliosis: a subject-specific musculoskeletal modelling study

**DOI:** 10.64898/2026.10.27.762384

**Authors:** Rounak Bhattacharya, Bhavuk Garg, Rajesh Malhotra, Rajdeep Ghosh, Anoop Chawla, Kaushik Mukherjee

**Affiliations:** School of Interdisciplinary Research, Indian Institute of Technology Delhi, New Delhi 110 016, Delhi, India; Department of Orthopaedics, All India Institute of Medical Sciences, New Delhi 110029, Delhi, India; Currently at Department of Orthopaedics, Max Super Speciality Hospital, Saket, New Delhi 110 017, Delhi, India; Currently at Department of Orthopaedics, Indraprastha Apollo Hospital, New Delhi 110029, Delhi, India; Department of Mechanical Engineering, Indian Institute of Technology Delhi, New Delhi 110 016, Delhi, India; Currently at Department of Mechanical Engineering, National Institute of Technology Agartala, Agartala 799 046, Tripura, India

**Keywords:** Adolescent idiopathic scoliosis, Load carrying, Musculoskeletal modelling, Intervertebral loading, Paraspinal muscle forces

## Abstract

Load carrying is a common activity during adolescence, yet its biomechanical consequences for the structurally asymmetric spine in adolescent idiopathic scoliosis (AIS) remain poorly understood. This study investigated how load magnitude, carrying height, and side of application influence spinal loading and paraspinal muscle forces in AIS. A subject-specific musculoskeletal model of a 16-year-old female with a 59° right-convex thoracic curve was used to simulate bilateral and unilateral load carrying at the hand, chest, and above-shoulder levels, with loads corresponding to 5%, 10%, and 15% of body weight. Intervertebral compressive and lateral forces and erector spinae (ES) and multifidus (MF) muscle forces were evaluated. Spinal loading increased progressively with load magnitude and carrying height. At 15% body weight, compressive forces at the curve apex increased from 511–566 N at hand level to 721–831 N above the shoulder. Unilateral carrying produced distinct side-dependent responses: convex-side carrying generated greater compressive forces, whereas concave-side carrying generated greater lateral forces, which were concentrated around the curve apex. Paraspinal muscles also exhibited muscle-specific responses. ES forces primarily reflected the imposed external asymmetry, with unilateral carrying eliciting predominantly contralateral recruitment, whereas MF maintained a characteristic convex-side dominance concentrated around the curve apex. These findings demonstrate that load carrying in AIS produces a coupled biomechanical response governed by both the external loading configuration and the pre-existing structural asymmetry of the spine, highlighting the importance of considering loading direction and muscular adaptation alongside load magnitude when evaluating everyday carrying activities in AIS.

## 1. Introduction

Adolescent idiopathic scoliosis (AIS) is a complex spinal disorder defined by a lateral spinal curvature (exceeding 10° Cobb angle) in the coronal plane. The three-dimensional deformity can vary considerably in curve location, magnitude, and pattern (Cheng et al., 2015; Choudhry et al., 2016). A higher prevalence is observed in females, particularly with larger and progressive curves. Despite extensive clinical and biomechanical investigations, the mechanisms underlying the onset and progression of AIS remain incompletely understood (Fadzan and Bettany-Saltikov, 2017; Grauers et al., 2016; Kikanloo et al., 2019). As a result, there remains considerable interest in understanding how everyday physical demands may influence the biomechanical response of the scoliotic spine.

Load-carrying is a common functional activity during adolescence and imposes additional mechanical demands on the spinal column. The biomechanical response to external loading depends not only on the magnitude of the load, but also on the manner in which it is carried, thereby influencing spinal loading and the paraspinal muscle recruitment required to maintain postural stability (McGill et al., 2013; Antonius Rohlmann et al., 2014; A. Rohlmann et al., 2014; Rose et al., 2013). In individuals with AIS, load-carrying has been associated with back pain and has been suggested to influence deformity progression (Chow et al., 2006; Sahli et al., 2013). Nevertheless, the biomechanical response of the scoliotic spine to externally applied loads remains poorly understood.

Direct measurement of intervertebral loading or paraspinal muscle forces is challenging in adolescents, as available *in vivo* techniques are invasive and have limited applicability in this population. Musculoskeletal (MSK) modelling provides a non-invasive computational approach for estimating internal joint loading and muscle forces that are difficult to measure *in vivo* (Bruno et al., 2017; Katelyn A. Burkhart et al., 2018; Dreischarf et al., 2016). MSK models have been used extensively to investigate lifting and load-carrying tasks in healthy populations (Akhavanfar et al., 2024; Banks et al., 2023; Beaucage-Gauvreau et al., 2019; Kim and Zhang, 2017). However, these findings cannot be assumed to represent individuals with AIS, in whom altered spinal curvature changes vertebral alignment and muscle moment arms.

To the authors’ knowledge, previous investigations of load-carrying in AIS have focussed primarily on backpack carriage (Schmid et al., 2020b), while the biomechanical consequences of other common carrying strategies remain largely unexplored. In particular, little is known about how different forms of hand-held load carrying influence spinal loading in AIS. In addition, unilateral load carrying is of particular interest because placing the load on one side of the body introduces an external mechanical asymmetry that may interact with the pre-existing asymmetry of the scoliotic spine. This is clinically relevant in the context of the “vicious cycle” hypothesis (Stokes et al., 1996), whereby asymmetric mechanical loading may promote asymmetric vertebral growth and further aggravate the spinal deformity (Makino et al., 2016). A systematic evaluation of these load-carrying conditions may therefore provide further insight into the mechanical response of the spine in AIS.

Therefore, the present study aimed to investigate the effects of load-carrying on spinal loading and paraspinal muscle forces in an individual with AIS using a subject-specific MSK modelling approach. Load-carrying was simulated at the hand, chest, and above-shoulder levels under various load magnitudes. Both bilateral and unilateral carrying conditions were evaluated, with unilateral loads applied on either the concave or convex side of the curve. Intervertebral compressive and lateral forces were quantified across the scoliotic curve, together with changes in erector spinae (ES) and multifidus (MF) muscle forces. This framework enabled a systematic evaluation of how load-carrying height, carrying mode, and load magnitude collectively influence the biomechanical response of the scoliotic spine.

## 2. Methods

### 2.1. Study participant

The present study was based on a 16-year-old female with AIS (height: 1.59 m; weight: 55 kg), whose radiographic data were collected in a previous study (Bhattacharya et al., 2026). The patient presented with a right-convex thoracic curve (Cobb angle: 59°), with the curve apex located at T10. Spatially calibrated biplanar radiographs (EOS Imaging, Paris, France), which allow simultaneous acquisition of frontal and lateral images, were obtained in the upright standing posture. Ethical approval was granted by the Institutional Review Board of the All-India Institute of Medical Sciences, New Delhi (IEC-248/04.03.2022), and written informed consent was obtained from the patient’s guardians.

### 2.2. Subject-specific MSK model

A subject-specific AIS model was developed from an age- and sex-matched OpenSim full-body musculoskeletal model of an adolescent (Schmid et al., 2020a). The detailed model development procedure has been described in our previous study (Bhattacharya et al., 2026). Briefly, subject-specific spinal alignment was reconstructed from the biplanar radiographs by identifying anatomical landmarks on each vertebra. These landmarks were used to determine vertebral positions and orientations in the frontal and sagittal planes, as well as the intervertebral joint locations (Bassani et al., 2017). The reconstructed alignment was subsequently incorporated into the scaled MSK model to reproduce the patient-specific spinal deformity. The rib cage and upper-limb segments were then realigned to maintain their anatomical configuration, while the head–neck segment was adjusted to preserve a horizontal gaze in the neutral standing posture (Schmid et al., 2020b). Finally, the rib joints were locked, and the intercostal muscles were excluded to reduce the computational complexity of the model (Rauber et al., 2024).

**Fig. 1.**
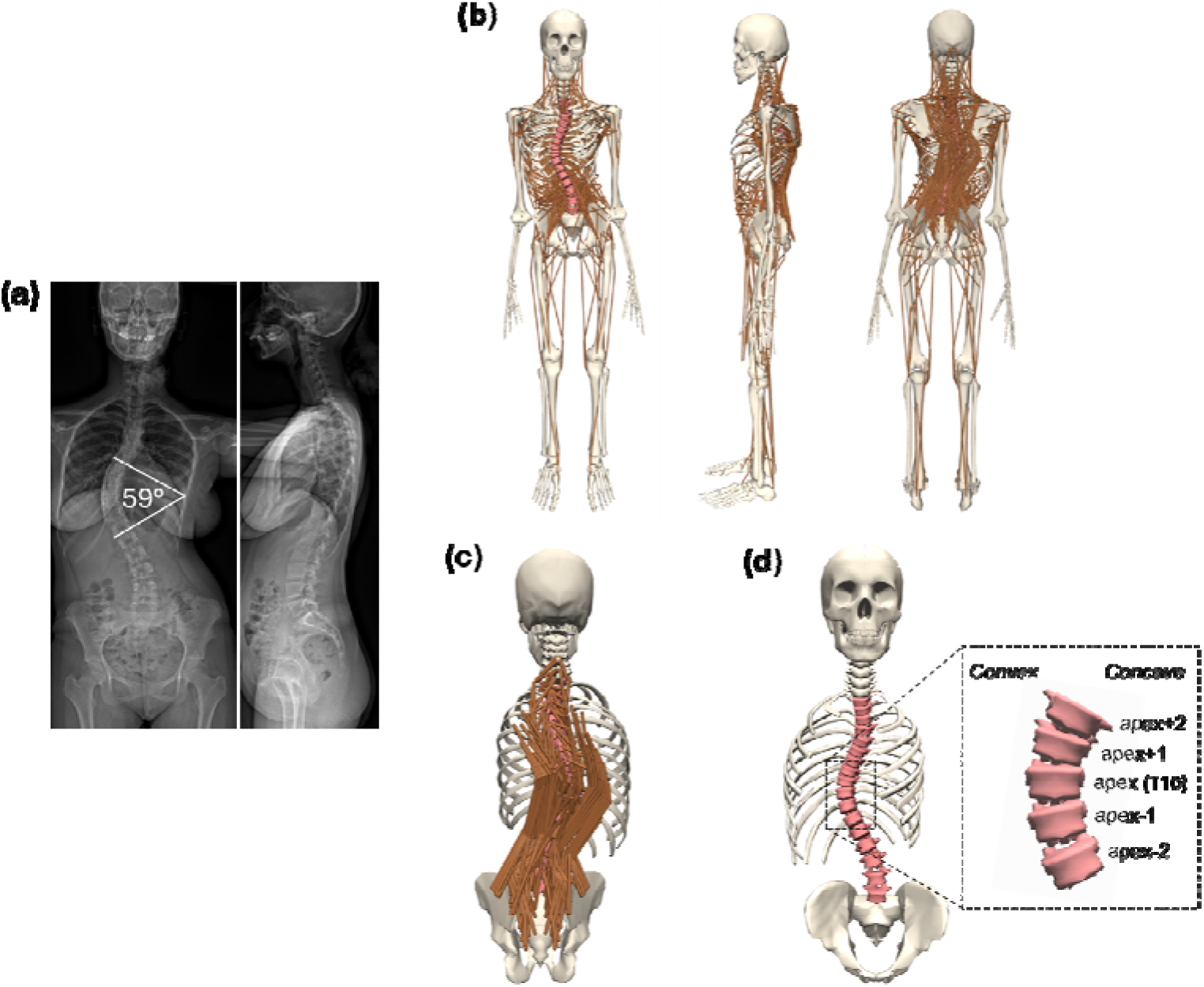
Subject-specific MSK model of the participant with AIS. (a) Biplanar radiographs; (b) full-body MSK model; (c) paraspinal musculature; and (d) spinal geometry showing the curve apex (T10), adjacent levels, and the convex and concave sides of the curve.

The model was previously evaluated against reported *in vivo* (Kwok et al., 2015) and MSK-derived (Barba et al., 2021; Schmid et al., 2020b) muscle activation patterns in individuals with AIS, as described in our previous study (Bhattacharya et al., 2026). In brief, predicted erector spinae activation was assessed using convex-to-concave activation ratios in the thoracic and lumbar regions and at spinal levels surrounding the curve apex. The predicted activation patterns showed good agreement with previously reported *in vivo* measurements (Kwok et al., 2015) and MSK modelling predictions (Barba et al., 2021; Schmid et al., 2020b).

### 2.3. Simulation of load-carrying conditions

Three load-carrying conditions were simulated: hand-level, chest-level, and above-shoulder carrying. For hand-level carrying, the arms were positioned alongside the body with the elbows extended. For chest-level carrying, the elbows were flexed to position the hands in front of the chest, whereas for above-shoulder carrying, the shoulders were flexed to position the hands above shoulder level. For each carrying condition, three load-carrying modes were simulated: bilateral, concave-side, and convex-side carrying. During bilateral carrying, the load was equally distributed between the two hands, whereas during unilateral carrying, it was applied to either the concave or convex side of the hand. External loads corresponding to 5%, 10%, and 15% of the patient’s body weight (BW) were applied vertically downward to the hand segments. An unloaded neutral standing condition was also simulated to serve as a reference for comparison.

**Fig. 2.**
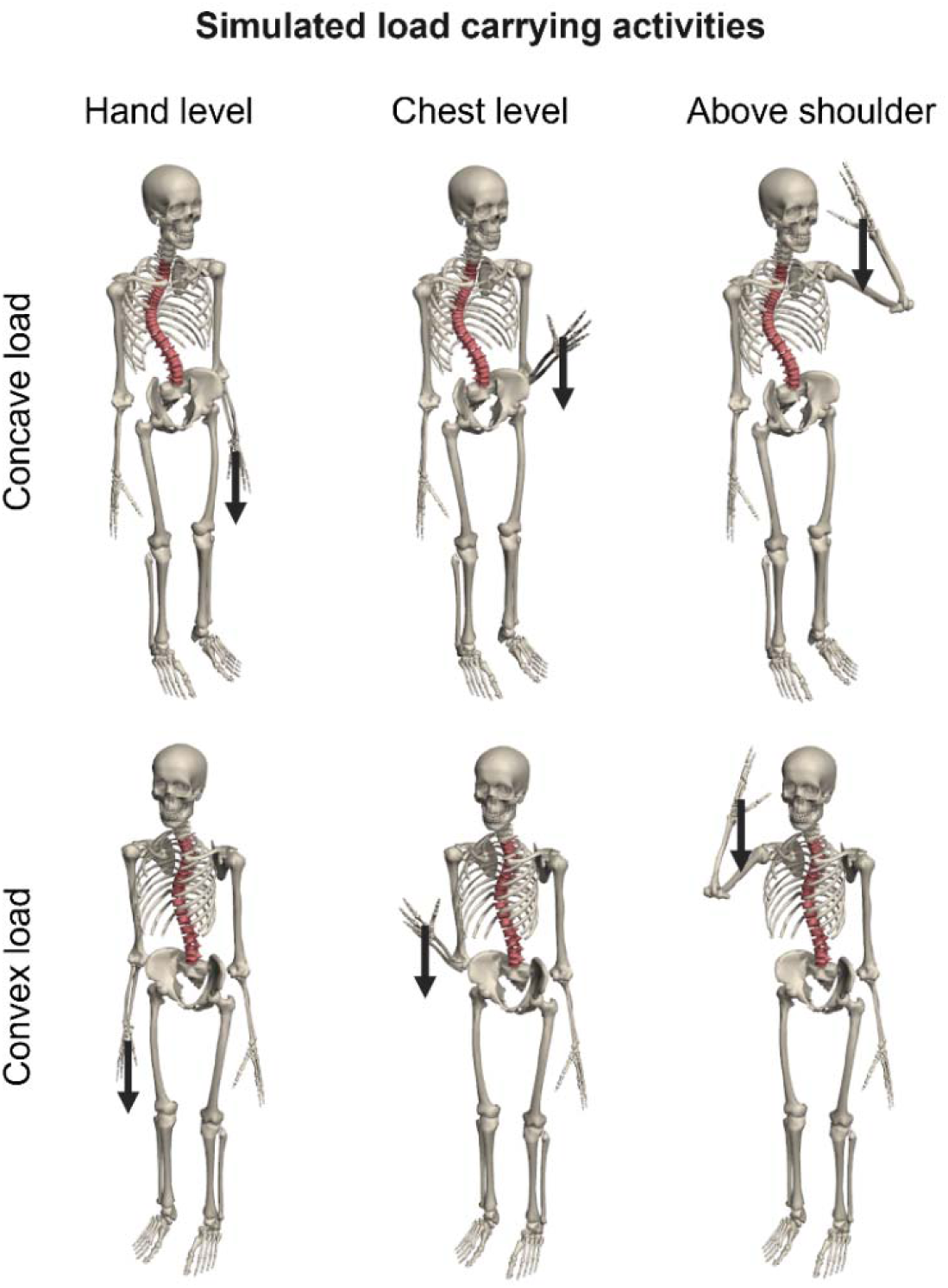
Simulated load-carrying conditions at hand, chest, and above-shoulder levels. For illustrative clarity, only the unilateral concave- and convex-side carrying conditions are shown.

### 2.4. Musculoskeletal simulation

An inverse simulation pipeline (Seth et al., 2018) was used to estimate muscle forces and intervertebral reaction forces across the simulated load-carrying conditions. Muscle forces were predicted using static optimisation, with the objective of minimising the sum of squared muscle activations (Erdemir et al., 2007). Residual actuators were applied between the calcaneus and the ground to provide external mechanical support during the simulations (Katelyn A Burkhart et al., 2018). Joint reaction analysis was subsequently used to determine the intervertebral reaction forces.

Intervertebral compressive and lateral forces were evaluated at the curve apex and at two vertebral levels above and below the apex. These levels were selected to capture the regional distribution of spinal loading around the curve apex. Erector spinae (ES) and multifidus (MF) muscle forces were evaluated separately on the concave and convex sides of the curve. Total muscle force on each side was obtained by summing the forces generated by the corresponding muscle fascicles from T1 to L5. Level-wise muscle forces were also determined from the fascicles crossing each vertebral mid-plane (Bruno et al., 2015). Muscle force imbalance was quantified as the difference between the convex- and concave-side forces (convex − concave), with positive values indicating convex-side dominance and negative values indicating concave-side dominance.

## 3. Results

### 3.1. Effect of load carrying on intervertebral reaction forces

Compressive forces increased with load-carrying height, with substantially greater forces observed during chest- and above-shoulder-level carrying than during hand-level carrying (Fig. 3). At 15% BW, the compressive force at the curve apex ranged from 511–566 N at hand level, increasing to 709–796 N at chest level and 721–831 N at above-shoulder level. Carrying mode further influenced compressive loading, with convex-side carrying consistently producing greater forces than concave-side carrying. At 15% BW, the apex compressive forces for concave-/convex-side carrying were 516/566 N at hand level, 722/796 N at chest level, and 758/831 N at above-shoulder level. The distribution of compressive forces around the curve apex also varied with load-carrying height, with a relatively uniform distribution across the evaluated levels during hand-level carrying and greater forces at the lower evaluated levels during chest- and above-shoulder-level carrying.

**Fig. 3.**
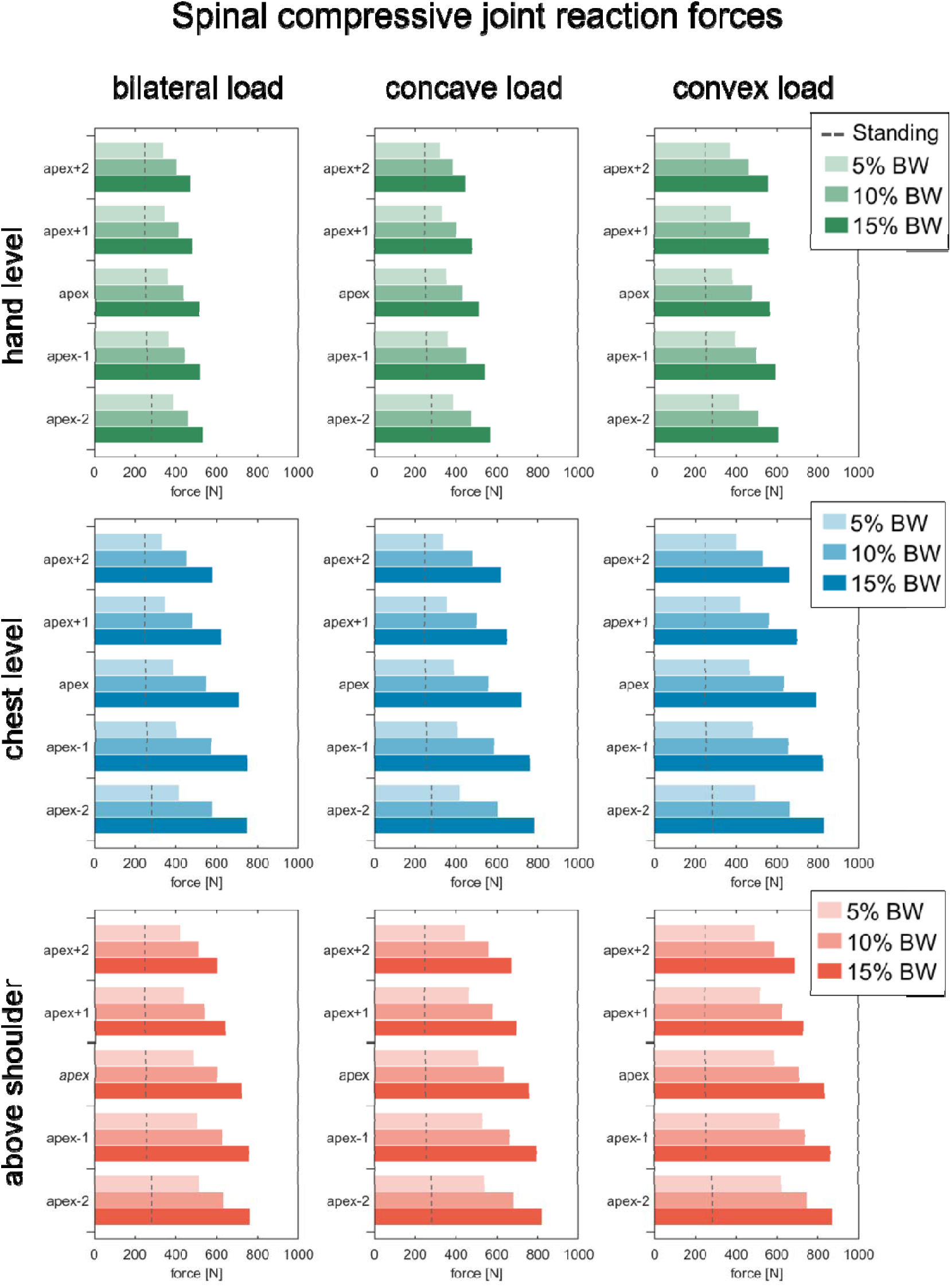
Spinal compressive forces at the curve apex and at two levels above and below during simulated load-carrying activities. Bars with progressively deeper shades indicate increasing load magnitudes (5%, 10%, and 15% BW). The dashed vertical line denotes the reference condition of neutral standing without load.

In contrast to compressive loading, lateral forces showed a distinct response to carrying mode (Fig. 4), with concave-side carrying consistently producing greater lateral forces than convex-side carrying. At 15% BW, the apex lateral forces for concave-/convex-side carrying were 203/165 N at hand level, 279/242 N at chest level, and 298/260 N at above-shoulder level. Load-carrying height also affected lateral loading. At 15% BW, the lateral force at the curve apex increased from 149–203 N at hand level to 221–279 N at chest level and 236–298 N at above-shoulder level. Across all load-carrying heights and modes, lateral forces were predominantly concentrated around the curve apex. Both compressive and lateral forces increased consistently as load magnitude increased from 5% to 15% BW.

**Fig. 4.**
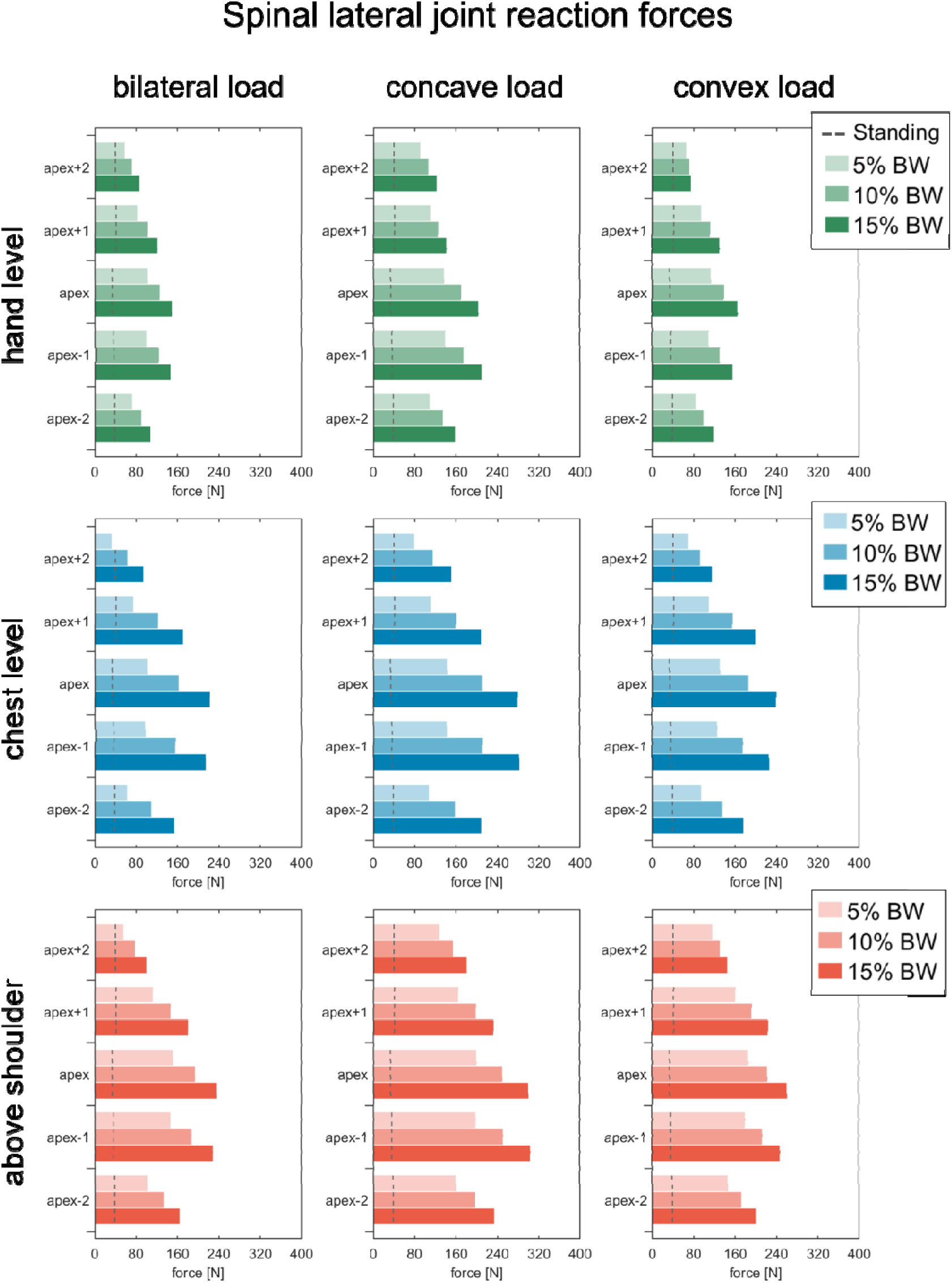
Spinal lateral forces at the curve apex and at two levels above and below during simulated load-carrying activities. Bars with progressively deeper shades indicate increasing load magnitudes (5%, 10%, and 15% BW). The dashed vertical line denotes the reference condition of neutral standing without load.

### 3.2. Effect of load carrying on paraspinal muscle forces

Erector spinae (ES) muscle forces were strongly influenced by carrying mode (Fig. 5a), with unilateral carrying producing a predominantly contralateral muscle response. Concave-side carrying increased convex-side ES forces, whereas convex-side carrying increased concave-side ES forces. At 15% BW during concave-side carrying, concave/convex ES forces were 117/275 N at hand level, 287/455 N at chest level, and 258/422 N at above-shoulder level. During convex-side carrying, the corresponding values were 224/149 N, 384/300 N, and 353/272 N, respectively.

**Fig. 5.**
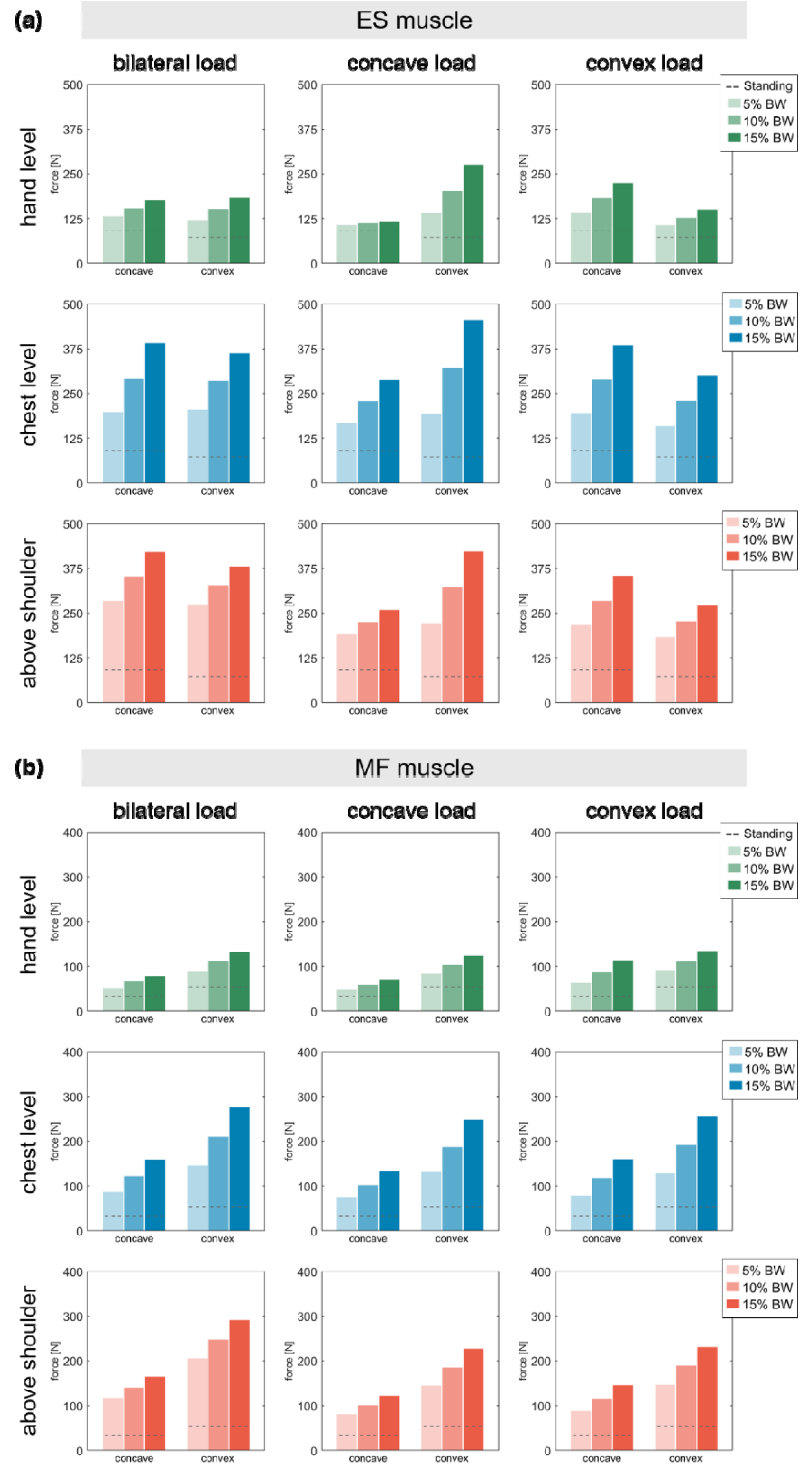
(a) Erector spinae and (b) multifidus muscle forces on the concave and convex sides during simulated load-carrying activities. Bars represent 5%, 10%, and 15% BW loads; dashed lines indicate neutral standing without load.

Multifidus (MF) muscle forces exhibited a more consistent pattern of convex-side dominance across the different carrying modes (Fig. 5b). MF forces increased with both load-carrying height and load magnitude, with greater forces generally observed during chest- and above-shoulder-level carrying than during hand-level carrying. At 15% BW during bilateral carrying, concave/convex MF forces were 78/132 N at hand level, 158/275 N at chest level, and 164/290 N at above-shoulder level. A similar height-dependent increase was observed during unilateral carrying.

Level-wise analysis showed distinct distributions of muscle force imbalance between the ES and MF muscles. ES muscle force imbalance was predominantly concentrated in the lumbar region (Fig. 6a), with greater imbalance observed at higher load magnitudes during unilateral carrying. In contrast, MF muscle force imbalance was primarily concentrated around the curve apex (Fig. 6b), with predominantly convex-side dominance across the simulated load-carrying conditions.

**Fig. 6.**
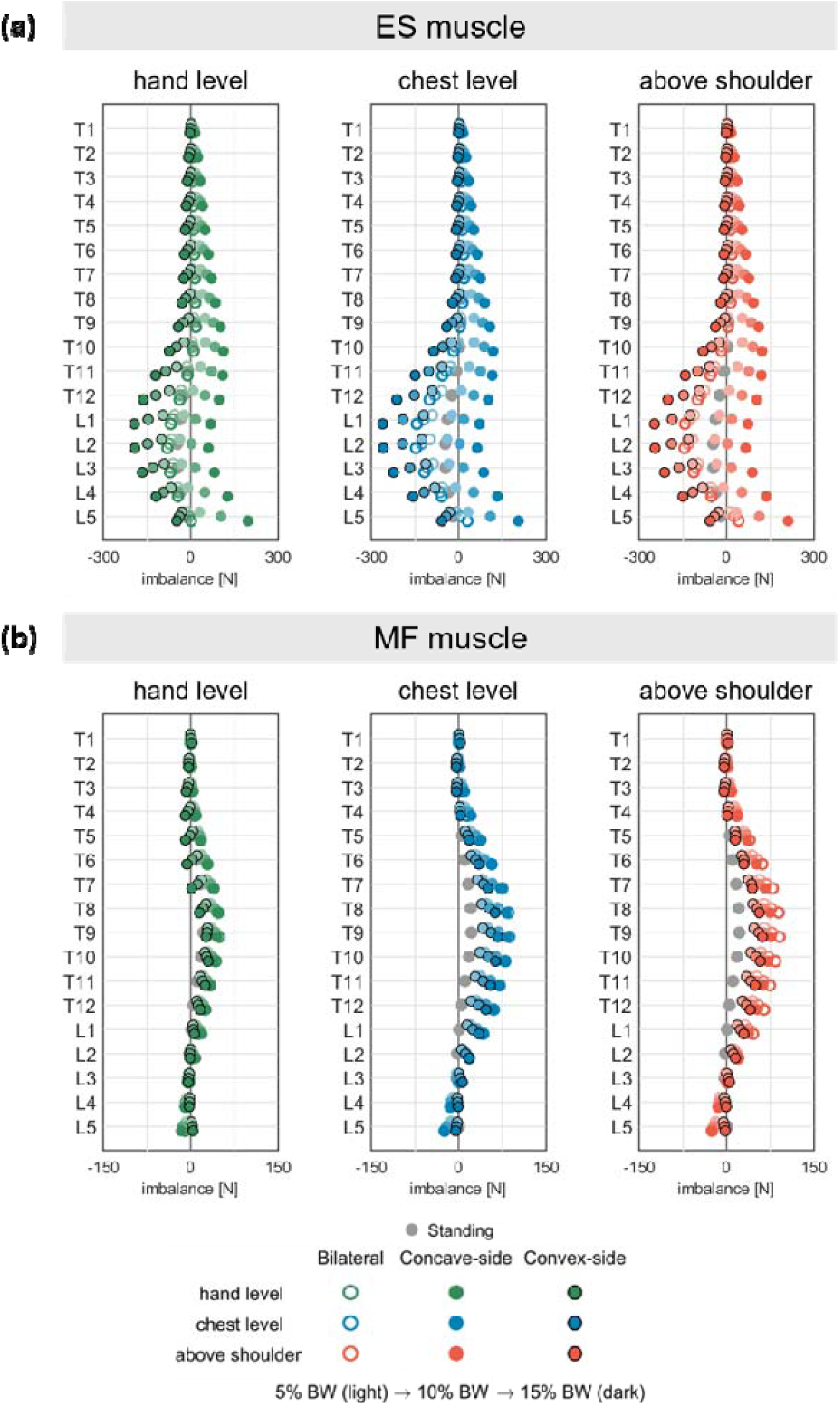
Muscle force imbalance for (a) erector spinae (ES) and (b) multifidus (MF) across spinal levels during simulated load-carrying activities. Imbalance was calculated as (convex – concave); thus, negative and positive values indicate concave- and convex-side dominance, respectively.

## 4. Discussion

This study investigated the biomechanical effects of different load-carrying tasks on spinal joint reaction forces and paraspinal muscle forces in an AIS patient using a subject-specific MSK model. The results demonstrated that spinal loading was strongly influenced by the load-carrying configuration, with intervertebral reaction forces increasing progressively with increasing load magnitude and carrying height. Moreover, load-carrying in different modes, i.e., unilateral or bilateral carrying, also altered the distribution of paraspinal muscle forces, reflecting the muscular adjustments required to maintain spinal equilibrium under varying external loading conditions. Together, these findings demonstrate how variations in load-carrying configuration influence spinal loading and paraspinal muscle stabilisation demand in AIS.

Load-carrying height substantially influenced compressive forces around the scoliotic curve apex. Compressive forces increased progressively from hand-level to chest- and above-shoulder-level carrying, potentially reflecting greater muscular demands required to maintain static equilibrium. Although compressive forces generally increased toward the lower evaluated spinal levels, consistent with the cumulative transmission of spinal loading (Han et al., 2013; Meszaros-Beller et al., 2023), this difference became more pronounced at elevated carrying positions. Furthermore, increasing load magnitude amplified compressive forces across all evaluated levels, indicating that both carrying height and load magnitude contribute to the mechanical demands imposed on the scoliotic spine.

In addition to carrying height, the carrying mode also influenced compressive loading. Convex-side carrying produced the highest compressive forces, followed by concave-side carrying and bilateral carrying. This pattern indicates that unilateral load carrying interacts with the pre-existing spinal asymmetry, resulting in a direction-dependent compressive response. Convex-side carrying likely aligns the external load more directly with the deformed spinal axis, promoting greater axial force transmission across the curve. These findings are consistent with Schmid et al. (2020b), who reported greater compressive loading during convex-side backpack carrying than concave-side carrying in individuals with AIS under upright standing conditions. Together, these findings suggest that the mechanical consequences of unilateral carrying in AIS depend not only on load magnitude and position, but also on the side on which the load is carried.

In contrast to compressive forces, lateral shear forces showed a different trend, with concave-side carrying generating the highest lateral forces, followed by convex-side carrying and bilateral carrying. The higher lateral forces observed during concave-side carrying may reflect altered spinal load transmission, whereby a greater proportion of the applied external load is transmitted laterally rather than vertically due to the deformed spinal geometry. This altered load transfer is further accentuated around the curve apex, where the geometric asymmetry of the scoliotic spine is greatest. Importantly, the opposite side-dependent responses observed for compressive and lateral forces indicate that the mechanical effect of unilateral carrying cannot be characterised by compression alone. The importance of considering lateral shear forces in scoliosis has also been highlighted by Barba et al. (2021), who showed that neglecting lateral shear forces and considering only compressive loading may provide an incomplete representation of spinal loading in AIS. The present findings further highlight the importance of considering multiple components of intervertebral loading when evaluating the biomechanical response of the scoliotic spine. Future studies should further investigate the role of lateral shear loading in AIS, particularly its potential contribution to structural adaptation and curve progression.

The observed patterns of paraspinal muscle forces suggest that load-carrying substantially influences trunk muscle recruitment strategies in AIS. Even under symmetric bilateral loading, ES muscle forces showed a slight concave-convex asymmetry, reflecting the underlying structural asymmetry of the scoliotic spine. During unilateral carrying, the ES muscle showed a clear compensatory response, with greater forces on the opposite side of the carried load to counterbalance the external load. Similar responses have been reported in healthy individuals (Kim and Zhang, 2017; Motmans et al., 2006; Zhao et al., 2023). Using MSK modelling, Kim and Zhang (2017) reported increased paraspinal muscle forces during asymmetric lifting, while Zhao et al. (2023) observed greater contralateral paraspinal muscle forces during unilateral handbag carrying. However, in the present AIS model, the muscle force distribution differed depending on the side of load application, with concave-side carrying producing a greater ES muscle imbalance than convex-side carrying. This suggests that the ES response depends not only on unilateral loading but also on the specific side of the curve on which the load is carried, reflecting the interaction between the applied load and the pre-existing spinal asymmetry in AIS.

In contrast to the load-dependent response of the ES muscle, the MF muscle showed a more consistent pattern of convex-side dominance across the simulated carrying conditions. In healthy individuals, greater contralateral lumbar multifidus (MF) activation has been reported during unilateral hand-held loading, suggesting that MF recruitment is influenced by the side of load application (Paek et al., 2022). In comparison, the persistent convex-side MF dominance observed in the present study is consistent with Schmid et al. (2020b), who reported greater MF activation on the convex side around the curve apex in AIS. This persistence of MF asymmetry despite changes in carrying mode suggests that its force distribution may be more strongly associated with the underlying spinal geometry than with the direction of the external load. The level-wise analysis further showed that MF imbalance was predominantly concentrated around the curve apex, where the structural asymmetry of the scoliotic spine is most pronounced. This localisation may reflect the role of the MF in providing segmental stabilisation across the deformed spinal segments. Thus, the different responses of the ES and MF suggest distinct functional adaptations to load carrying, with the ES responding more dynamically to external load asymmetry, while the MF maintains a more persistent curvature-associated asymmetry.

While the present results provide important biomechanical insight into load-dependent spinal responses in AIS, they do not provide a definitive clinical conclusion regarding whether bilateral or unilateral load carrying is more beneficial for minimising curve progression or preventing complications such as joint degeneration or back pain. Although convex-side carrying resulted in higher compressive forces and may therefore increase the mechanical burden on the spine, its long-term influence on curve progression remains uncertain. According to the Hueter– Volkmann principle, bone growth is slowed under compression and accelerated under distraction (De Sèze and Cugy, 2012). Therefore, the increased compressive loading on specific regions of the vertebrae could potentially modulate growth patterns and may influence vertebral development in scoliosis. However, whether such effects would be corrective or detrimental cannot be inferred from the present findings. The observed changes in lateral forces add further complexity to this interpretation. Increased lateral shear, particularly during concave-side carrying, amplified the pre-existing asymmetric load distribution across the scoliotic curvature and may influence structural remodelling near the curve apex. However, the biological consequences of lateral shear in the scoliotic spine, particularly its effects on tissue adaptation and vertebral growth, remain poorly understood. Therefore, caution is warranted in translating these findings directly into clinical recommendations regarding optimal load-carrying strategies in AIS.

Several limitations should be considered when interpreting the present findings. First, the model did not account for patient-specific skeletal morphology; incorporating CT/MRI-derived anatomy may improve the predictive accuracy of the model. Second, muscle forces were estimated using static optimisation, which assumes an optimal pattern of muscle recruitment; however, this may not fully capture the neuromuscular strategies adopted in AIS, where compensatory or non-optimal recruitment patterns may influence muscle force distribution. Third, the simulated conditions did not account for postural adaptations that may occur during real-life load carrying. Future studies incorporating motion-capture data could better represent these compensatory responses and provide a more realistic assessment of spinal loading. Finally, the analysis was based on a single subject with a specific curve configuration. The findings may therefore not be representative of the broader AIS population, given the substantial variability in curve pattern, severity, and spinal morphology.

## 5. Conclusions

The present study demonstrates that load carrying substantially alters the mechanical environment of the scoliotic spine, with greater spinal loading observed at higher load magnitudes and elevated carrying positions. Unilateral carrying further produced distinct side-dependent responses, with convex-side carrying generating greater compressive forces and concave-side carrying generating greater lateral forces. These loading patterns were accompanied by muscle-specific adaptations, with the erector spinae responding primarily to the imposed external asymmetry, whereas the multifidus retained a curvature-associated force imbalance concentrated around the curve apex. Collectively, these findings indicate that the mechanical consequences of load-carrying in AIS are determined not simply by the magnitude of the applied load, but by the interplay between carrying position, side of application, and the pre-existing structural asymmetry of the spine. Further studies across different curve patterns and real-life load-carrying scenarios are needed to determine whether these acute biomechanical responses contribute to long-term spinal adaptation or curve progression.

## Acknowledgement and Funding

The authors are grateful to the Indian Institute of Technology Delhi and the All-India Institute of Medical Sciences, New Delhi, for providing the facilities and infrastructure required to conduct this study.

## CRediT authorship contribution statement

**Rounak Bhattacharya:** Data Curation, Formal Analysis, Investigation, Methodology, Software, Validation, Visualisation, Writing – original draft. **Bhavuk Garg:** Conceptualisation, Methodology, Patient Recruitment Supervision, Writing – review & editing. **Rajesh Malhotra:** Conceptualisation, Methodology, Patient Recruitment Supervision, Writing – review & editing. **Rajdeep Ghosh:** Investigation, Methodology, Writing – review & editing. **Anoop Chawla:** Conceptualisation, Methodology, Resources, Supervision, Writing – review & editing. **Kaushik Mukherjee:** Conceptualisation, Methodology, Resources, Supervision, Writing – review & editing.

## Declaration of competing interest

There are no possible conflicts of interest for the authors in terms of funding support, research, authorship, or publishing of this paper.

## Statements of ethical approval

Ethical approval was obtained from the Institute Ethics Committee (IEC) of the All-India Institute of Medical Sciences, New Delhi, India, under approval number IEC-248/04.03.2022.

## Notes

### Competing Interest Statement

The authors have declared no competing interest.

